# USP22 controls type III interferon signaling and SARS-CoV-2 infection through activation of STING

**DOI:** 10.1101/2022.02.01.478628

**Authors:** Rebekka Karlowitz, Megan L. Stanifer, Jens Roedig, Geoffroy Andrieux, Denisa Bojkova, Sonja Smith, Lisa Kowald, Ralf Schubert, Melanie Boerries, Jindrich Cinatl, Steeve Boulant, Sjoerd J. L. van Wijk

**Author notes:** Corresponding author: Sjoerd J. L. van Wijk, Institute for Experimental Cancer Research in Pediatrics, Goethe University Frankfurt, Komturstrasse 3a, 60528 Frankfurt am Main, Germany.

## Abstract

Pattern recognition receptors (PRRs) and interferons (IFNs) serve as essential antiviral defense against SARS-CoV-2, the causative agent of the COVID-19 pandemic. Type III IFN (IFN-λ) exhibit cell-type specific and long-lasting functions in autoinflammation, tumorigenesis and antiviral defense. Here, we identify the deubiquitinating enzyme USP22 as central regulator of basal IFN-λ secretion and SARS-CoV-2 infections in native human intestinal epithelial cells (hIECs). USP22-deficient hIECs strongly upregulate genes involved in IFN signaling and viral defense, including numerous IFN-stimulated genes (ISGs), with increased secretion of IFN-λ and enhanced STAT1 signaling, even in the absence of exogenous IFNs or viral infection. Interestingly, USP22 controls basal and 2’3’-cGAMP-induced STING activation and loss of STING reversed STAT activation and ISG and IFN-λ expression. Intriguingly, USP22-deficient hIECs are protected against SARS-CoV-2 infection, viral replication and the formation of *de novo* infectious particles, in a STING-dependent manner. These findings reveal USP22 as central host regulator of STING and type III IFN signaling, with important implications for SARS-CoV-2 infection and antiviral defense.

## Introduction

Sensing of “non-self” is a key feature of innate immunity and underlies the recognition of viruses, bacteria and fungi, but also plays important roles in cancer and auto-immune diseases ^1, 2^. Pattern recognition receptors (PRRs), like Toll-like receptors (TLRs), Nucleotide-binding oligomerization domain (NOD)-like receptors (NLRs) and retinoic acid-inducible gene 1 protein (RIG-I)-like receptors (RLRs) are essential components of innate immune signaling and selectively recognize pathogen-associated molecular patterns (PAMPs). Dedicated PRRs, like TLR3, RIG-I, Melanoma differentiation-associated protein 5 (MDA5) and Cyclic GMP-AMP synthase (cGAS)-Stimulator of interferon genes protein (STING) recognize viral double stranded RNA (dsRNA) and dsDNA, and are important sensors for infections with RNA and DNA viruses, as well as infections with retroviruses ^1–3^. Whereas TLR3 recognizes dsRNA in endosomes, the prototypical RLRs, RIG-I and MDA5, sense cytosolic dsRNAs, while cGAS-STING detects viral dsDNA ^1–4^. STING is activated either directly via viral dsDNA, through the STING agonist 2’3’-cGAMP generated by the cyclic GMP-AMP synthase cGAS upon detection of viral dsDNA, or indirectly via RIG-1 and MDA5 ^5^. Activated STING interacts with TANK-binding kinase 1 (TBK1) and activates interferon regulatory factor (IRF) 1, −3, and −7 and Nuclear factor-κB (NF-κB), leading to the initiation of anti-viral and inflammatory transcriptional programs, including interferon-stimulated genes (ISGs) and interferons (IFN) ^5–8^.

Interferons (IFNs) are secreted cytokines with important roles in immunity and anti-viral responses. IFN signaling relies on Janus kinase-Signal transducer and activator of transcription (JAK-STAT) activation, phosphorylation of STAT1/2 and the induction of ISG and IFN gene expression that influence viral replication ^9, 10^. Although the vast majority of cell types can be triggered to express type I (IFN-α, -β, -ε, -κ and -ω) and type III (IFN-λ1, -λ2, -λ3 and -λ4) IFNs, the expression of IFN-specific receptors is cell type restricted and determines IFN responses. For example, the type I IFN receptor (IFNAR) is ubiquitously expressed in many tissues, whereas expression of the type III IFN receptor IFNLR1 is mainly limited to epithelial cells, e.g. the gastro-intestinal and respiratory epithelium ^6–8, 11, 12^. Although type I and type III IFNs induce similar ISG signatures, type I IFNs generally trigger a more rapid increase and decay of ISG expression ^7^. In addition, IFN-λs have been described to be first-in-line against viral infections and might inhibit viral spread without triggering inflammatory responses, depending on IFN-λ receptor expression ^7, 13, 14^.

The novel severe acute respiratory syndrome coronavirus 2 (SARS-CoV-2) is the causative agent of the pandemic Coronavirus disease 2019 (COVID-19) and belongs to the human coronaviruses (HCoV) that also includes SARS-CoV and MERS-CoV ^15^. In many patients with severe COVID-19, SARS-CoV-2 infection induces the secretion of highly pro-inflammatory cytokines through cGAS-STING and NF-κB-mediated signaling ^16, 17^. Type I and III IFNs are important regulators of host viral defense against SARS-CoV-2 ^6–8, 11, 12, 18, 19^, but at the same time, SARS-CoV-2 evades immune recognition via IFN and ISG suppression ^10, 20^. Prolonged expression of low basal levels of type I and III IFNs might prime host responses against virus infection, including SARS-CoV-2 ^21–24^. Although type III IFNs restrict SARS-CoV-2 infection in intestinal and airway epithelial cells ^19, 25–29^ and STING agonism reduces SARS-CoV-2 infection ^30–33^, context-dependent damaging effects of type III IFNs on airway epithelia during viral infections have been described as well ^34, 35^.

Innate immunity, PRRs and IFN signaling is closely regulated by ubiquitination, both by the host machinery as well as through viral E3 ligases and deubiquitinating enzymes (DUBs) that hijack the host ubiquitin machinery ^36^. STING, RIG-I, TLR3 and TBK1 are positively and negatively regulated by differential modification of polyubiquitin chains, including K11, K27, K48 and K63 linked chains ^37, 38^, by a variety of E3 ligases, such as TRIM56 ^39^, TRIM32 ^40^, MUL1 ^41^, AMFR ^42^, RNF5 ^43^ and TRIM29 ^44^ and RNF26 ^45^. The interplay and functional consequences of ubiquitin modifications are complex and include proteasomal degradation as well as stabilization of protein-protein interactions. Importantly, IFN and anti-viral signaling are also heavily regulated by DUBs, like USP13 ^46^, USP35 ^47^ and CYLD ^48^

Ubiquitin-specific peptidase 22 (USP22) is a DUB that is part of the deubiquitination module of the Spt-Ada-Gcn5-acetyltransferase (SAGA) complex, through which it regulates transcription via the control of histone H2A K119 and H2B K120 monoubiquitination (H2AK119ub1 and H2BK120ub1, respectively) ^49–51^. Recently, additional USP22 substrates have emerged, with important roles in cell fate regulation and programmed cell death ^52–54^. Interestingly, USP22 has mostly been associated with IFN signaling and ISG expression upon infection with viruses ^55, 56^. However, up till now, the mechanisms how USP22 primes PRR and IFN signaling and prepares against anti-viral defense in native, uninfected settings remains unknown.

In light of the current COVID-19 pandemic and potential future pathogenic coronaviruses, identifying host factors that control SARS-CoV-2 infection is of extreme relevance. The roles of type III IFN in SARS-CoV-2 infections are only starting to emerge and are determined by tissue-specific factors as well. Here, we are the first to identify USP22 as a negative regulator of basal ISG expression, JAK/STAT activation and IFN signaling, even in the absence of exogenous IFNs or viral infection. Our findings elucidate USP22 as crucial host factor in shaping SARS-CoV-2 antiviral defense by priming cellular anti-viral responsiveness prior to virus infection. Loss of USP22 in native, human intestinal epithelial cells (hIECs) triggers a strong upregulation of ISGs and, specifically, the type III IFN IFN-λ, mediated by STING. USP22 controls basal and 2’3’-cGAMP-induced STING ubiquitination, phosphorylation and activation, and combined loss of USP22 and STING rescues ISG expression, STAT signaling and IFN-λ production. Importantly, we found that USP22-deficient hIECs are prominently protected against SARS-CoV-2 infection, replication and the formation of novel infectious viral particles, which can be partially reversed by loss of STING expression.

## Results

### Profiling USP22-mediated gene expression in HT-29 hIECs

Substrate-specific deubiquitination is a central determinant of ubiquitin homeostasis and regulates receptor activation and internalization, proteasomal degradation and transcription. For the ubiquitin-specific protease USP22, both transcriptional and extranuclear targets have been identified. As part of the DUB module of the SAGA complex, USP22 regulates transcriptional elongation via H2AK119ub1 and H2BK120ub1 ^49–51^. Up till now, the spectrum of target genes regulated by USP22 largely remains unclear, partially due to organism-, cell-and context-dependent redundancy in alternative DUBs that might compensate for loss of USP22 ^57^. We previously reported that CRISPR/Cas9-mediated knockout (KO) of USP22 in human colon carcinoma cell line HT-29 affects RIPK3 ubiquitination during necroptosis, but without inducing major changes in RIPK1, RIPK3 and MLKL gene expression ^54^, suggesting gene-specific regulation of USP22. To identify the spectrum of USP22-regulated genes, we profiled USP22-dependent changes in gene expression in the human intestinal epithelial cell (hIEC) line HT-29. Quantification of alterations in gene expression in two independent HT-29 USP22 KO single cell clones revealed a marked alteration in gene expression, with 401 genes up-regulated and 182 down-regulated (Figure 1A **and** Supplemental Figure 1A). Loss of USP22 expression was accompanied with changes in H2Bub1, but not H2Aub1 (Supplemental Figure 1B & C). Among the top-50 differentially regulated genes, 30 were up-and 20 downregulated, with an adjusted P-value of < 0.05 (Figure 1B). Genes upregulated in both USP22 KO clones (#16 and #62) compared to control (non-human target: NHT) HT-29 cells include genes that encode for proteins involved in growth and differentiation, like Transforming Growth Factor β-1 (TGFB1), Tumor-associated calcium signal transducer 2 (TACSTD2) and Tyrosine-protein kinase Mer (MERTK) and the cytosolic RNA- and DNA sensor DExD/H-Box Helicase 60 (DDX60). Downregulated genes include USP22, mitochondrial adenylate kinase 4 (AK4) that is involved in the regulation of mitochondrial function and ATP production ^58^ and regenerating islet-derived protein 4 (REG4), a carbohydrate-binding lectin that has been identified as marker for deep crypt secretory cells (DSCs) that acts as niche for Lgr5-positive stem cells in the colon ^59^. Differential regulation of gene expression, as well as loss of USP22 expression, was also demonstrated by independent qRT-PCR of the USP22-dependent upregulated genes TGFB1, SLFN5, TGM2 and DDX60, as well as downregulation of USP22, CXCR4 and AK4 (Figure 1C), confirming the quality of the microarray.

**Figure 1:**
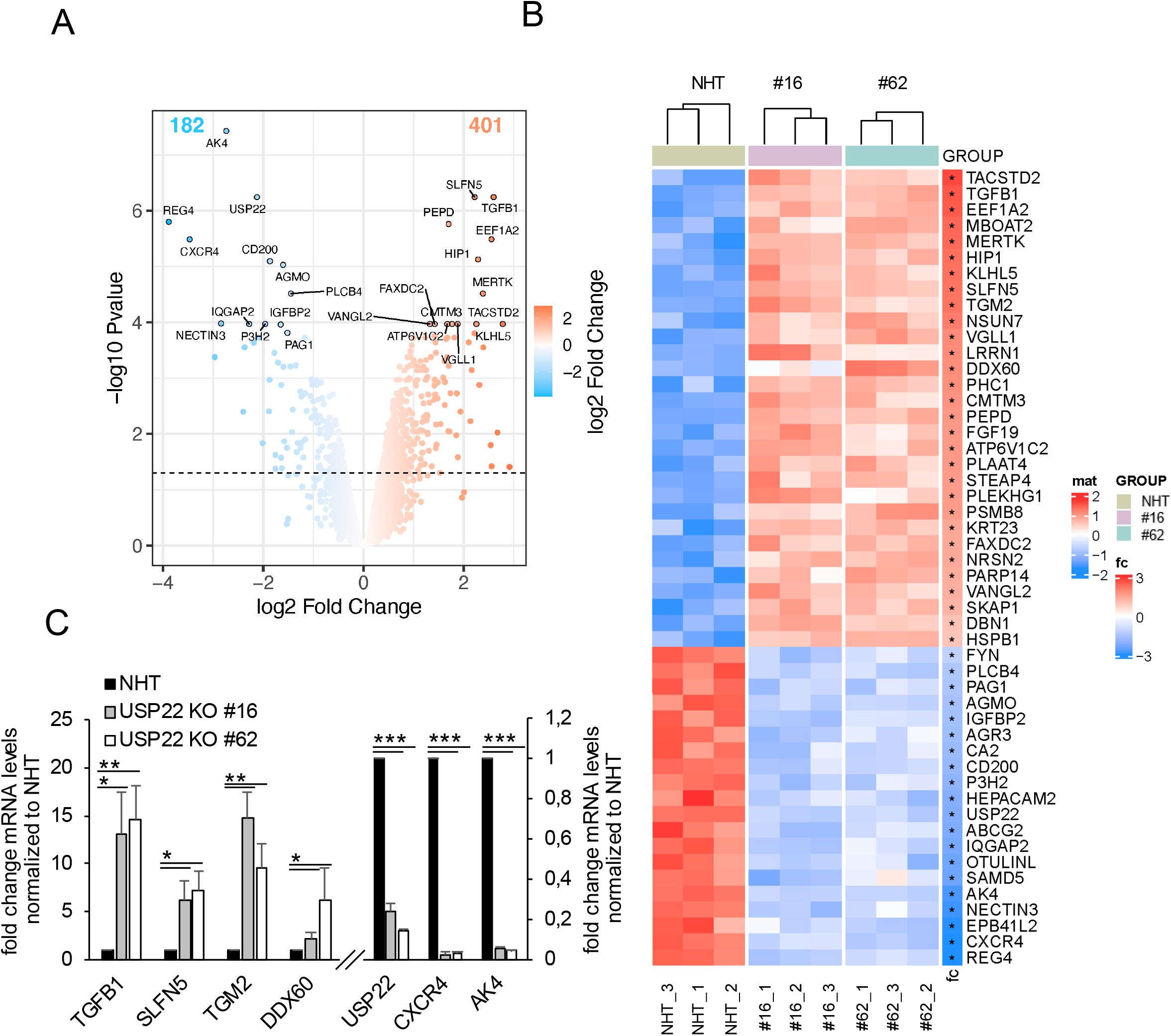
Profiling USP22-mediated gene expression in HT-29 hIECs. **A.** Volcano plot showing the differential gene expression patterns of two independent single-cell HT-29 USP22 CRISPR/Cas9 KO clones (#16 and #62) compared to CRISPR/Cas9 control (NHT) HT-29 cells. Color code represents the log2 foldchange compared to NHT. **B.** Heatmap of the top-50 differentially regulated genes between HT-29 USP22 KO single clones #16 and #62 and the NHT control. Color coding represents the row-wise scaled (Z-score) RNA intensities. Genes are sorted according to their log2 fold change, compared to NHT. **C.** Basal mRNA expression levels of the indicated genes were determined in control and two independent USP22 KO HT-29 single clones using qRT-PCR. Gene expression was normalized against 28S mRNA and is presented as x-fold mRNA expression compared to NHT. Mean and SD of three independent experiments in triplicate are shown. *P < 0.05; **P < 0.01, ***P < 0.001.

### Loss of USP22 specifically enriches for genes involved in interferon signaling and response to viral infection

Next, gene-set enrichment analysis was performed on USP22-regulated genes to investigate if certain gene sets from gene ontology (GO) are specifically regulated by USP22. Interestingly, GO analysis revealed an enrichment of genes linked to type I and II interferon (IFN) signaling, as well as regulation of viral genome replication and several other viral processes, such as the regulation of viral genome replication, response to virus, response to IFN-γ, IFN-γ mediated signaling pathway in USP22 KO HT-29 cells as compared to control NHT HT-29 cells (Figure 2A). Interestingly, the GO terms of genes that are strongly downregulated are enriched in mitochondrial translation and gene expression, ribosomal and ribonucleoprotein complex biogenesis and the processing of tRNA, rRNA and ncRNA (Figure 2A).

**Figure 2:**
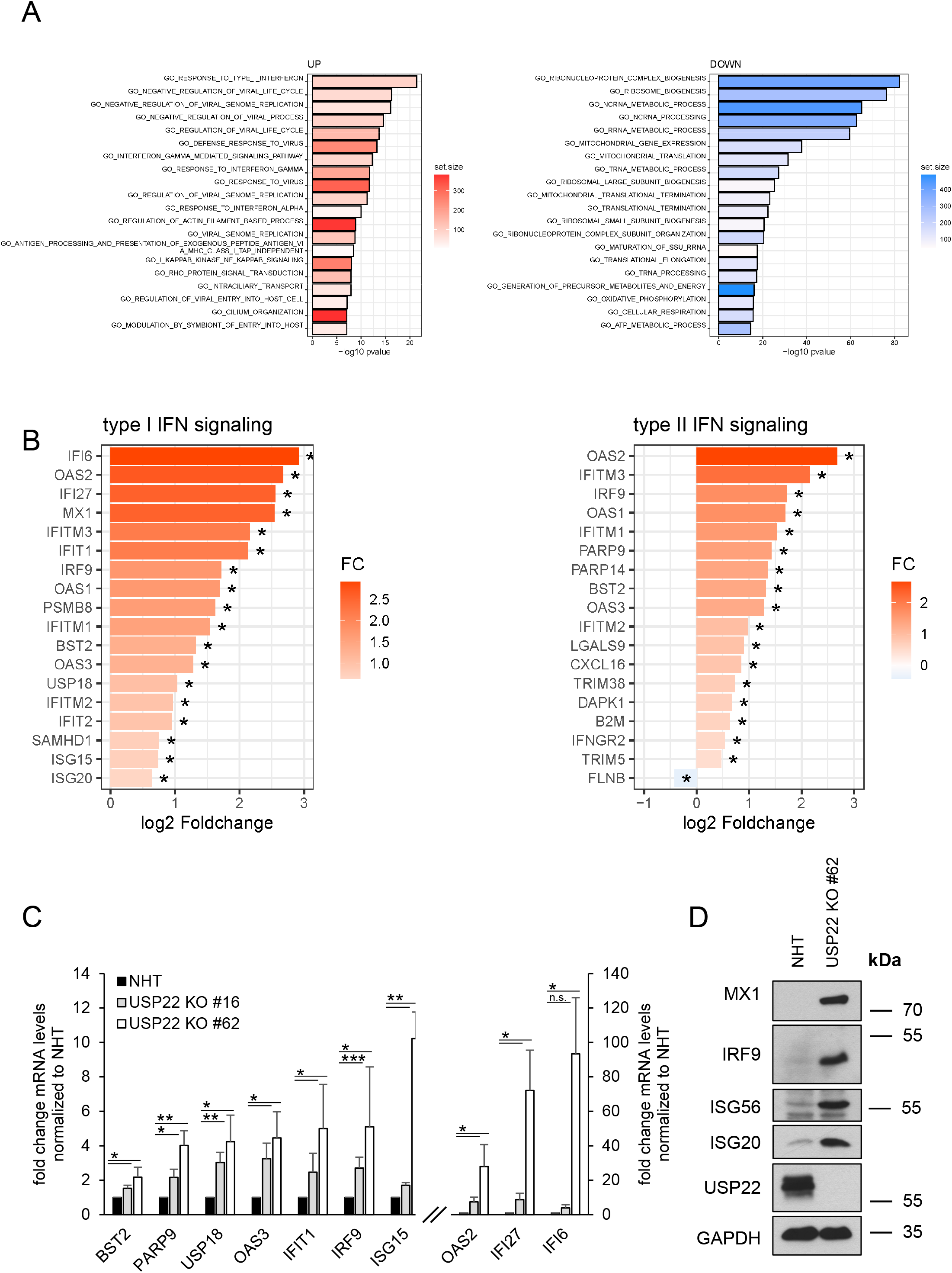
Loss of USP22 specifically enriches for genes involved in interferon signaling and response to viral infection. **A.** Bar plot showing the top-20 regulated Gene Ontology (GO) terms in two independent single-cell HT-29 USP22 CRISPR/Cas9 KO clones (#16 and #62) compared to control (non-human target: NHT) HT-29 cells. Color code represents the number of annotated genes within each gene set. **B.** Heatmap of differentially expressed genes contributing to the GO terms *response to type I interferon* (left) and *interferon gamma mediated signaling pathway* (right). Color code represents the log2 foldchange compared to NHT. **C.** Basal mRNA expression levels of GO enriched genes related to IFN signaling in control (NHT) and two independent USP22 KO HT-29 single clones using qRT-PCR. Gene expression was normalized against 28S mRNA and is presented as x-fold mRNA expression compared to NHT. Mean and SD of three independent experiments in triplicate are shown. *P < 0.05; **P < 0.01, ***P < 0.001, n.s. not significant. **D.** Western blot analysis of basal MX1, IRF9, ISG56, ISG20 and USP22 expression levels in control and USP22 KO HT-29 cells (clone USP22 KO #62). GAPDH served as loading control. Representative blots of at least two different independent experiments are shown.

Since previous studies suggest controversial roles of USP22 in IFN signaling ^55, 56, 60, 61^, we decided to further study USP22-dependent changes in genes involved in type I or type II IFN signaling (Figure 2B). Loss of USP22 leads to the upregulation of many IFN stimulated genes (ISGs), some with important functions in viral defense, like OAS1, −2 and −3, MX1 and IFI27, suggesting a potential role of USP22 in the regulation of interferon signaling and viral responses (Figure 2B). Among the upregulated genes were components of the ISGylation machinery, like the ubiquitin-like modifier ISG15 and the ISG15-specific DUB USP18 ^62^. To validate the USP22-regulated changes in gene expression, qRT-PCR confirmed the increased expression of several ISGs, like BST2, PARP9, USP18, OAS3, IFIT1, IRF9, ISG15, OAS2, IFI27 and IFI6 in two independent HT-29 USP22 KO clones (Figure 2C). In addition, increased protein expression of MX1, IRF9, ISG56 and ISG20 could also be confirmed upon loss of USP22 (Figure 2D). These findings suggest that USP22 specifically controls the expression of genes involved in IFN signaling and virus defense, even in the absence of exogenous IFN stimulation or virus infection.

### USP22 negatively regulates STAT1 signaling and IFN-λ1 expression

The expression of ISGs is typically induced upon activation of IFN signaling pathways during pathogen invasion or autoinflammatory disease and serves to control inflammation and other defensive mechanisms ^9^. Additionally, several IFNs are constitutively expressed at low levels as well ^63^ to prime and increase cellular responsiveness of IFN signaling upon activation by external stimuli ^23^. Interestingly, the expression levels of pan-IFN-α and IFN-β mRNA were upregulated upon loss of USP22, compared to non-human target control HT-29 (Figure 3A). This was accompanied by an increase in the expression of STAT1, an IFN-regulated ISG itself ^64^ as well as STAT1 phosphorylation, suggesting activation of IFN signaling pathways in USP22 KO HT-29 cells, compared to control cells (Figure 3B). Interestingly, in contrast to mRNA levels, in-depth analysis of USP22-mediated alterations in the secretion of IFNs and IFN-related cytokines revealed only low basal levels of secreted IFN-α and IFN-β, suggesting that these cytokines might only weakly contribute to the observed ISG signature (Figure 3C). Surprisingly, the secretion and expression of IFN-λ1, a type III IFN, was strongly upregulated in USP22 KO HT-29 cells compared to control cells (Figure 3C & D). In addition, loss of USP22 expression also induced elevated basal secretion of the pro-inflammatory cytokines CXCL10 and IL-8 and minor changes in the secretion of IFN-α2 and GM-CSF, compared to controls. (Figure 3C). These findings suggest that USP22 negatively regulates IFN-λ1 expression and ISG induction. Since type III IFN-induced target genes largely overlap with genes regulated by type I IFNs ^65, 66^, type III IFN is likely the main IFN contributing to the USP22-dependent induction of ISG expression and STAT1 activation.

**Figure 3:**
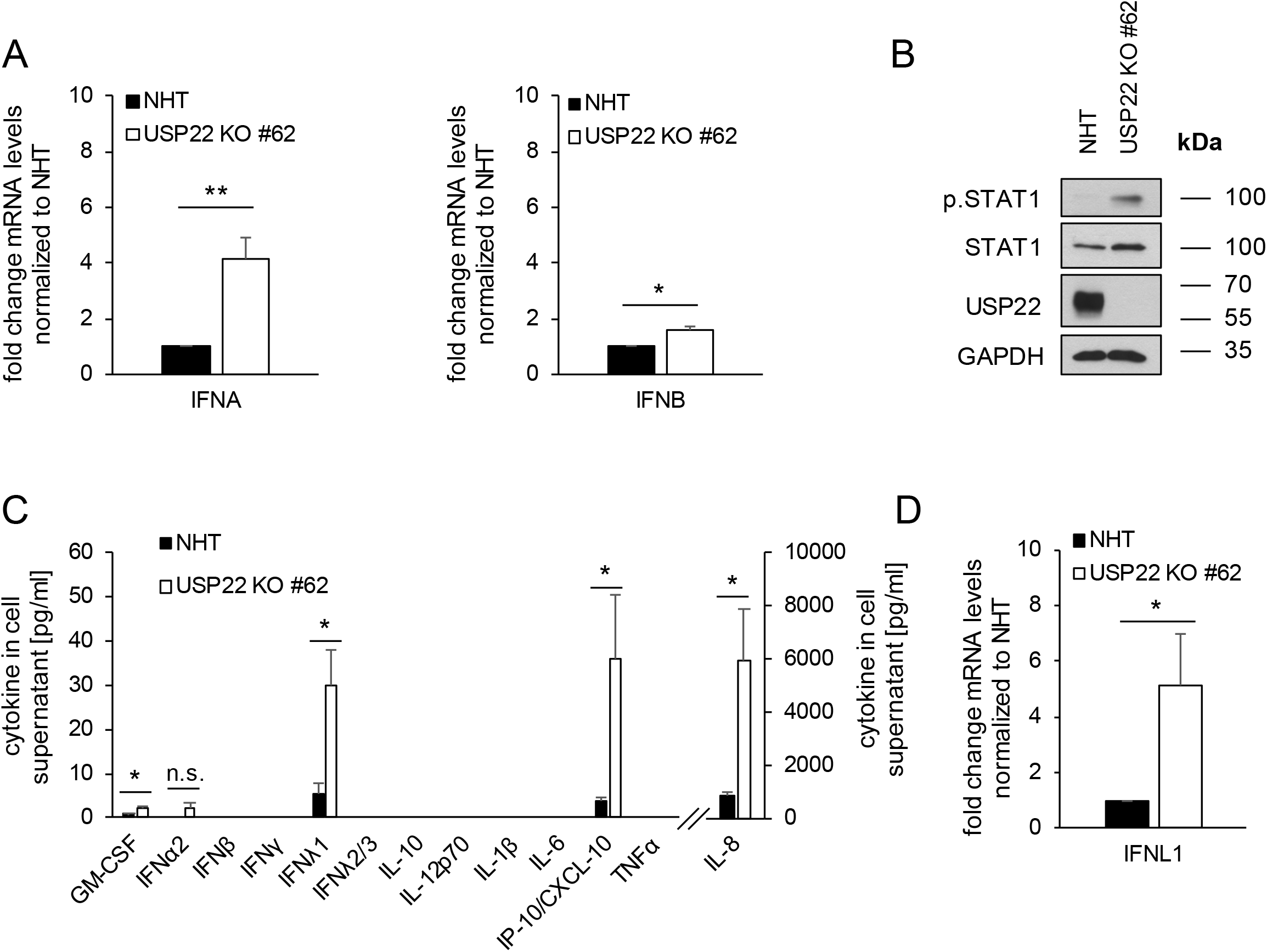
USP22 negatively regulates STAT1 signaling and IFN-λ1 expression. **A.** Basal mRNA expression levels of total IFNA (panIFNA) and IFNB1 in control (non-human target: NHT) and the CRISPR/Cas9-generated USP22 knock-out (KO) HT-29 single clone (USP22 KO #62). Gene expression was normalized against 28S mRNA and is presented as x-fold mRNA expression compared to NHT. Mean and SD of three independent experiments in triplicate are shown. *P < 0.05; **P < 0.01. **B.** Western blot analysis of basal phosphorylated and total levels of STAT1 and USP22 in control and USP22 KO HT-29 cells (USP22 KO #62). GAPDH served as loading control. Representative blots of at least two different independent experiments are shown. **C.** FACS-based analysis of the indicated basal secretion patterns of the viral defense cytokine panel in supernatants of control and USP22 KO HT-29 cells (USP22 KO #62). Data are presented as absolute levels of cytokines (in pg/ml). Samples below lower detection limit were set to zero, values above upper detection limit were set to detection limit. Mean and SD of three independent experiments in triplicate are shown. *P < 0.05; n.s. not significant. **D.** Basal mRNA expression levels of IFNL1 in control and USP22 KO HT-29 single clone (USP22 KO #62). Gene expression was normalized against 28S mRNA and is presented as x-fold mRNA expression compared to NHT. Mean and SD of three independent experiments in triplicate are shown. *P < 0.05.

### USP22 regulates type III IFN signaling via STING

Loss of USP22 expression specifically upregulates genes involved in IFN and viral responses. Within the context of viral infections, viral PAMPs, such as viral dsRNA and dsDNA are sensed by PRRs, like RIG-I, MDA5 and TLR3, and PRR activation mediates strong expression of IFNs and ISGs ^2, 5, 10^. Loss of USP22 leads to increased expression of RIG-I, MDA5 and TLR3 (Figure 4A). To investigate if these PRRs are functionally involved in USP22-mediated increased ISG signaling, the expression of RIG-I/DDX58, MDA5/IFIH1 and TLR3 was ablated with CRISPR/Cas9 in NHT and USP22 KO HT-29 cells (Figure 4B - D). Interestingly, despite efficient KO of the individual PRRs in both NHT and USP22 KO HT-29 cells, additional deletion of RIG-I, MDA5 or TLR3 did not decrease USP22-dependent STAT1 phosphorylation or ISG56 expression (Figure 4B - D). Interestingly, USP22-TLR3 double knockout (dKO) HT-29 cells even exhibit an increase in phosphorylated and total STAT1 levels as well as ISG56 expression, suggesting potential TLR3-specific effects of USP22 (Figure 4D).

**Figure 4:**
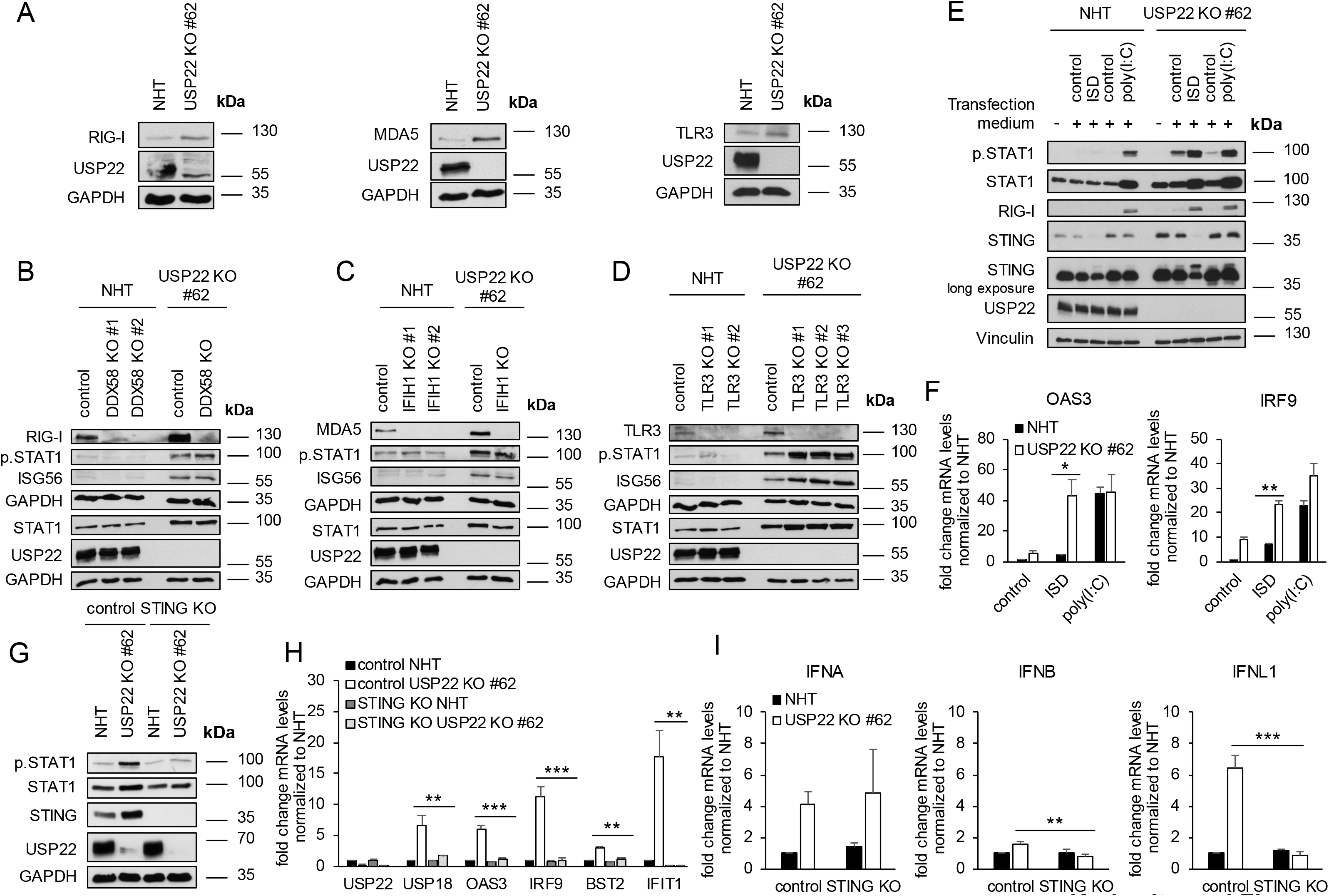
USP22 regulates type III IFN signaling via STING. **A.** Western blot analysis of basal RIG-I, MDA5, TLR3 and USP22 expression levels in control (non-human target: NHT) and the CRISPR/Cas9-generated USP22 knock-out (KO) HT-29 single clone (USP22 KO #62). GAPDH served as loading control. Representative blots of at least two different independent experiments are shown. **B.** Western blot analysis of basal RIG-I, phosphorylated and total STAT1, ISG56 and USP22 expression levels in control, USP22 KO HT-29 cells (USP22 KO #62) as well as two NHT-control and one USP22-DDX58 double KO (dKO) HT-29 single clones. GAPDH served as loading control. Representative blots of at least two different independent experiments are shown. **C.** Idem as ***B.***, one MDA5/IFIH1 KO single clone instead of RIG-I/DDX58. **D.** Idem as ***B.*** three TLR3 KO single clones instead of RIG-I/DDX58. **E.** Western blot analysis of phosphorylated and total STAT1, RIG-I, STING and USP22 expression levels in control and USP22 KO HT-29 cells (USP22 KO #62) subjected to transfection with transfection reagent (control) alone or in the presence of ISD and poly(I:C) (2µg/well) for 24 h. Vinculin served as loading control. Representative blots of at least two different independent experiments are shown. **F.** mRNA expression levels of OAS3 *(left)* and IRF9 *(right)* in control and USP22 knock-out (KO) HT-29 cells (USP22 KO #62) subjected to transfection with ISD and poly(I:C) (each 2 µg/well) for 24 h. Gene expression was normalized against 28S mRNA and is presented as x-fold mRNA expression compared to NHT. Mean and SD of three independent experiments in triplicate are shown. *P < 0.05; **P < 0.01; n.s. not significant. **G.** Western blot analysis of phosphorylated and total STAT1, STING and USP22 expression levels in control, USP22 KO HT-29 cells (USP22 KO #62) as well as in the indicated NHT, USP22, control and STING dKO cells. GAPDH served as loading control. Representative blots of at least two different independent experiments are shown. **H.** Basal mRNA expression levels of the indicated genes in control USP22 KO HT-29 cells (USP22 KO #62) as well as in the indicated NHT, USP22, control and STING dKO cells. Gene expression was normalized against 28S mRNA and is presented as x-fold mRNA expression compared to NHT. Mean and SD of three independent experiments in triplicate are shown. **P < 0.01; ***P < 0.001. **I.** Basal mRNA expression levels of IFNA, IFNB and IFNL1 in control USP22 KO HT-29 cells (USP22 KO #62) as well as in the indicated NHT, USP22, control and STING dKO cells. Gene expression was normalized against 28S mRNA and is presented as x-fold mRNA expression compared to NHT. Mean and SD of three independent experiments in triplicate are shown. **P < 0.01; ***P < 0.001.

An alternative source of IFN production might be from PRR-mediated detection of self-DNA (e.g. DNA damage and DNA double strand breaks), leading to the induction of IFN-α and IFN-λ via NF-κB signaling ^67^, as observed in several types of cancer. This is of special interest since USP22, apart from its role in transcriptional regulation, has also been associated with DNA damage responses ^68^ and V(D)J recombination and CSR *in vivo* by facilitating c-NHEJ ^69^. Since CRISPR/Cas9-mediated loss of USP22 did not lead to increased γH2AX levels in hIECs compared to controls (Supplemental Figure 2A) despite increased NF-κB signaling (Supplemental Figure 2B), it seems unlikely that DNA damage caused by loss of USP22 might contribute to IFN signaling.

Loss of RIG-I, MDA5 and TLR3 did not reverse the USP22-dependent IFN signature, suggesting that either functional redundancy between the selected PRRs could compensate for loss of individual PRRs or that additional PRRs are involved. Interestingly, expression of STING/TMEM173 was also increased in USP22 KO HT-29 cells (Figure 4E **and** Supplemental Figure 2C). STING can be activated via cGAS or indirectly via RIG-1 and MDA5, leading to complex formation with TBK1 and activation of IFN and NF-κB signaling ^5–8^. To further investigate the potential PRR redundancy and the involvement of STING, NHT and USP22 KO HT-29 cells were stimulated with the TLR3-, RIG-I- and MDA5-agonist polyinosinic:polycytidylic acid (poly(I:C)), or the 45-bp non-CpG oligomer IFN-stimulating DNA (ISD) from *Listeria monocytogenes* that strongly activates the STING-TBK1-IRF3 axis ^70, 71^. Intriguingly, whereas poly(I:C) induced a prominent increase in the levels of total and phosphorylated STAT1 in both NHT and USP22 KO cells, ISD selectively induced increases in total and phosphorylated STAT1 levels in USP22 KO cells, but not in NHT control cells, which was also reflected in a prominent ISD-mediated induction of RIG-I expression and STING activation (Figure 4E). In addition, ISD also induced strong expression of the representative ISGs OAS3 and IRF9 in USP22 KO cells compared to controls (Figure 4F). To confirm the role of STING in USP22-induced type III IFN signaling, USP22-STING dKO HT-29 cells were generated (Figure 4G). USP22-STING dKO cells exhibit strikingly reduced levels of basal and phosphorylated STAT1 protein compared to USP22 KO HT-29 cells (Figure 4G), suggesting a STING-dependent rescue of the USP22-dependent IFN signature. In line with this, USP22-induced ISG expression could be reversed as well in USP22-STING dKO HT-29 cells (Figure 4H). Additionally, USP22-mediated increases in IFN-λ expression could also largely be reduced upon USP22 STING dKO, whereas expression of IFN-α and IFN-β largely remains unaffected (Figure 4I). These findings reveal an important role of USP22 as negative regulator of STING-dependent type III IFN signaling in hIECs.

### USP22 negatively regulates STING activation and ubiquitination

The differential response to ISD, but not poly(I:C), and the reversal of the IFN signature in USP22-STING dKO hIECs suggests an important role of USP22 in the control of STING-induced type III IFN signaling. However, until now, the mechanisms of how USP22 regulates STING function remain unclear. Therefore, we subjected HT-29 control and USP22 KO cells to the STING agonist 2’3’-cGAMP and observed a fast, strong and more prolonged activation and phosphorylation of STING, as well as increased TBK1 and IRF3 phosphorylation (Figure 5A). In addition, the analysis of 2’3’-cGAMP-treated USP22 KO HT-29 cells revealed a very prominent increase in IFNL1 expression in USP22 KO cells, accompanied by increased IFNA and IFNB expression as well, but to a much lesser extent (Figure 5B).

**Figure 5:**
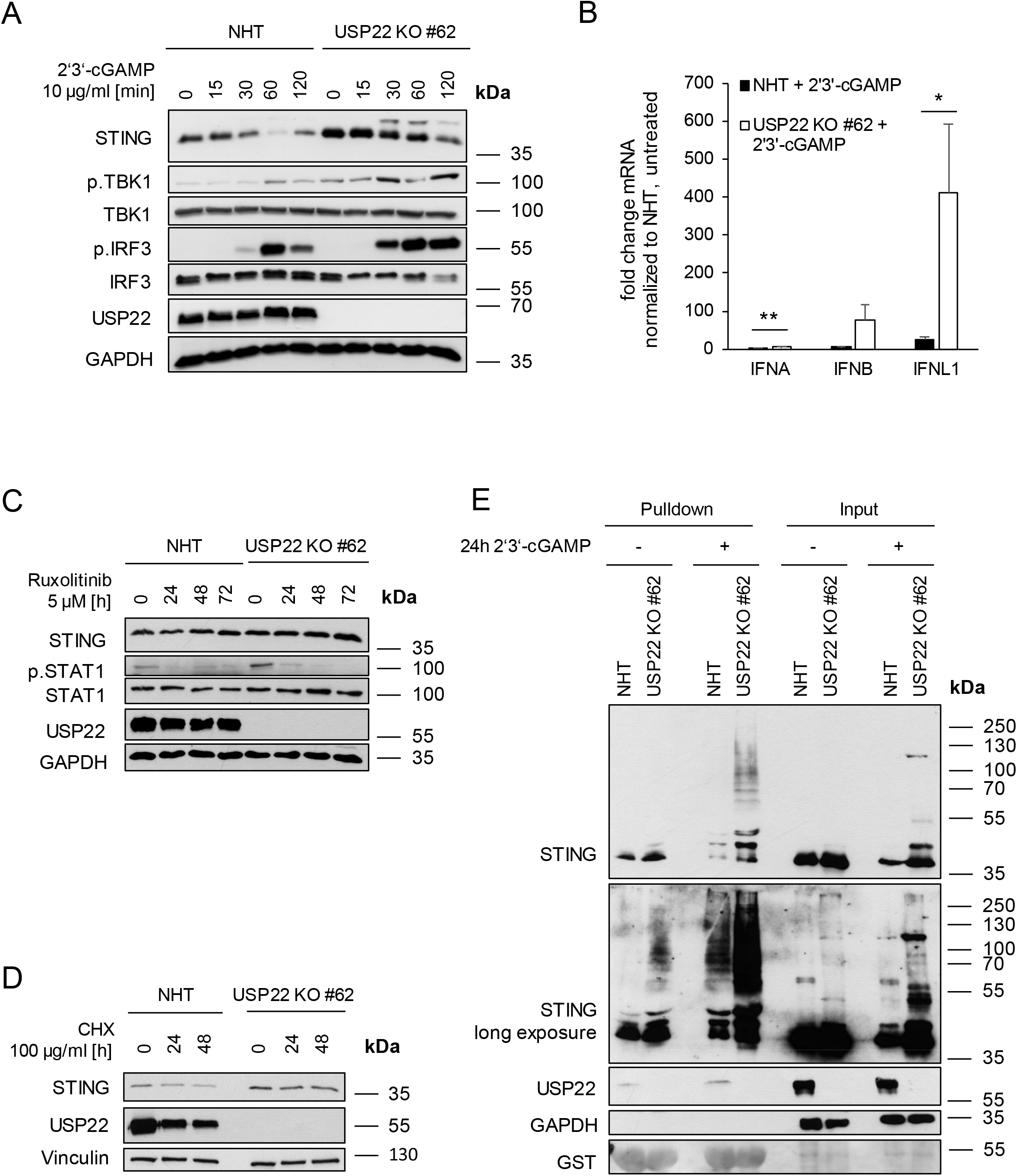
USP22 negatively regulates STING activation and ubiquitination. **A.** Western blot analysis of STING, phosphorylated and total TBK1, phosphorylated and total IRF3 and USP22 expression levels in control (non-human target: NHT) and CRISPR/Cas9-generated USP22 knock-out (KO) HT-29 single clone (USP22 KO #62) subjected to 2’3’-cGAMP (10 µg/ml) for the indicated timepoints. GAPDH served as loading control. Representative blots of at least two different independent experiments are shown. **B.** mRNA expression levels of IFNA, IFNB and IFNL1 in control and USP22 KO HT-29 cells (USP22 KO #62) subjected to 2’3’-cGAMP (10 µg/ml) for 3 h. Gene expression was normalized against 28S mRNA and is presented as x-fold mRNA expression compared to NHT. Mean and SD of three independent experiments in triplicate are shown. *P < 0.05; **P < 0.01. **C.** Western blot analysis of STING, phosphorylated and total STAT1 and USP22 expression levels in control and USP22 KO HT-29 cells (USP22 KO #62) subjected to the JAK/STAT inhibitor ruxolitinib (5 µM) for the indicated timepoints. GAPDH served as loading control. Representative blots of at least two different independent experiments are shown. **D.** Western blot analysis of STING and USP22 expression levels in control and USP22 KO HT-29 cells (USP22 KO #62) subjected to cycloheximide (CHX) (100 µg/ml) for the indicated timepoints. Vinculin served as loading control. Representative blots of at least two different independent experiments are shown. **E.** Western blot analysis of Tandem Ubiquitin Binding Entity (TUBE)-enriched ubiquitin-modified STING from control and USP22 KO HT-29 cells (USP22 KO #62) subjected to 2’3’-cGAMP (10 µg/ml) for 24 h. GAPDH served as loading control and Ponceau S staining confirms equal loading of GST-TUBE beads. Representative blots of at least two different independent experiments are shown.

Since STING expression is controlled by IFNs, constitutive IFN-mediated priming upon USP22 deficiency might underly the upregulation of STING. To investigate the relevance of auto- and paracrine IFN signaling in the regulation of STING expression, control and USP22 KO HT-29 cells were incubated with the JAK/STAT inhibitor ruxolitinib. JAK/STAT inhibition increased STING protein and mRNA expression levels in USP22 KO cells, compared to controls, suggesting that IFN-dependent auto- or paracrine activation of STING expression is unlikely (Figure 5C **and** Supplemental Figure 3A). Of note, USP22-mediated increases in STAT1 phosphorylation could be reversed with ruxolitinib (Figure 5C).

STING is reported to be modified with several types of ubiquitin chains that mediate context dependent effects, ranging from proteasomal degradation to the stimulation of signaling functions. STING protein levels were slightly stabilized in cycloheximide (CHX)-treated USP22 KO HT-29 cells compared to controls (Figure 5D). In line with these observations, basal and 2’3’-cGAMP-induced STING ubiquitination was also increased in USP22 KO HT-29 cells, compared to NHT control cells (Figure 5E). Together, these findings suggest that USP22-mediated effects on type III IFN might be predominantly regulated by activating STING ubiquitination and lesser through auto- or paracrine IFN signaling.

### Loss of USP22 protects against SARS-CoV-2 infection, replication and de novo infectious virus production in a STING-dependent manner

Previous studies revealed important, but highly context-dependent roles of STING and type III IFNs in the control of SARS-CoV-2 infection ^19, 30, 32, 33^. In addition, USP22 has been linked to viral signaling ^56^. To investigate the significance of USP22 and the resulting STING-mediated upregulation of type III IFN and ISG signaling for viral defense, the role of the USP22-STING axis was tested during SARS-CoV-2 infection. For this, we generated control and USP22 KO Caco-2 cells that express the virus receptors ACE-2 and TMPRSS2 and are susceptible to infection with SARS-CoV-2 virus ^19^. Loss of USP22 expression in Caco-2 cells triggered phosphorylation of STAT1 and increased expression of STING, compared to wild-type (WT) and NHT CRISPR/Cas9 control Caco-2 cells (Figure 6A). Increased USP22-dependent upregulation of IFN signaling in Caco-2 cells was also reflected in the increased expression of the antiviral ISGs IRF9 and OAS3 (Figure 6B). Intriguingly, USP22-deficient Caco-2 cells also express higher levels of IFN-λ1, compared to wild-type and CRISPR/Cas9 control non-human target cells, whereas IFN-α and IFN-β expression largely remained unaffected (Supplemental Figure 4A).

**Figure 6:**
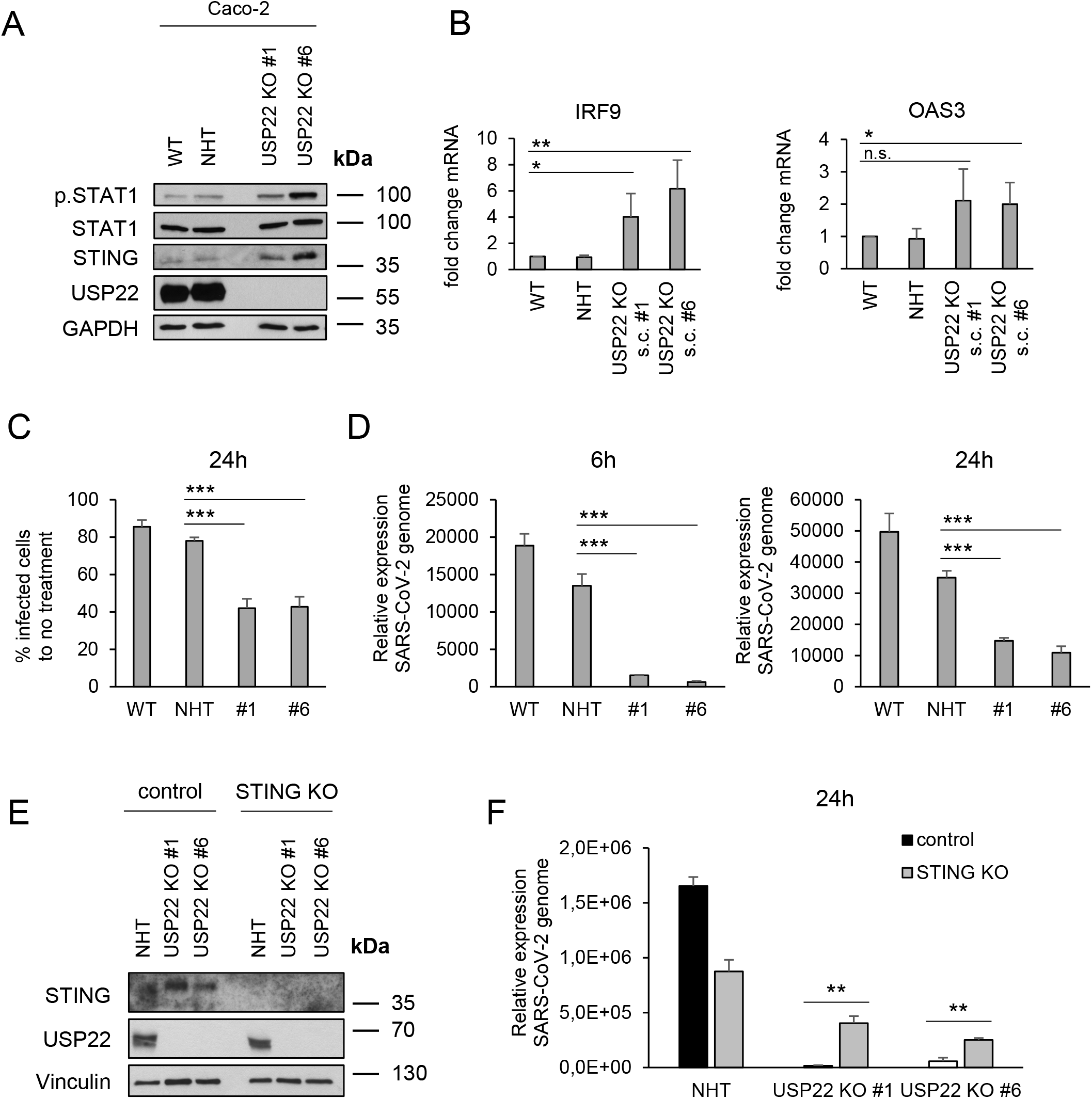
Loss of USP22 protects against SARS-CoV-2 infection, replication and de novo infectious virus production in a STING-dependent manner. **A.** Western blot analysis of phosphorylated and total STAT1, STING and USP22 expression levels in wild-type (WT), control (non-human target: NHT) and two CRISPR/Cas9-generated USP22 knock-out (KO) Caco-2 single clones (USP22 KO #1 and #6). GAPDH served as loading control. Representative blots of at least two different independent experiments are shown. **B.** Basal mRNA expression levels of IRF9 and OAS3 in WT, control and USP22 KO Caco-2 cells (USP22 KO #1 and #6). Gene expression was normalized against 28S mRNA and is presented as x-fold mRNA expression compared to NHT. Mean and SD of three independent experiments in triplicate are shown. *P < 0.05; **P < 0.01, n.s. not significant. **C.** Quantification of immunofluorescence-stained SARS-CoV-2-infected cells, normalized against non-infected cells. WT, control and USP22 KO Caco-2 cells (USP22 KO #1 and #6) were stained with anti-dsRNA (J2) at 24 hpi. Mean and SD of three independent experiments in triplicate are shown. ***P < 0.001. **D.** Quantification of relative SARS-CoV-2 genome expression of SARS-CoV-2-infected WT, control and USP22 KO Caco-2 cells (USP22 KO #1 and #6) at 6 hpi (left) and 24 hpi (right). Data are normalized against non-infected cells. Mean and SD of three independent experiments in triplicate are shown. ***P < 0.001. **E.** Western blot analysis of STING and USP22 expression levels in control-NHT, control-USP22 KO #1 and #6, STING-NHT and STING-USP22 KO #1 and #6 double KO (dKO) HT-29 cells. Vinculin served as loading control. Representative blots of at least two different independent experiments are shown. **F.** Quantification of relative SARS-CoV-2 genome expression of SARS-CoV-2-infected control-NHT, control-USP22 KO #1 and #6, STING-NHT and STING-USP22 KO #1 and #6 dKO HT-29 cells at 24 hpi. Mean and SD of three independent experiments in triplicate are shown. **P < 0.005.

To test the functional relevance of the increased antiviral signaling upon loss of USP22 expression, WT, NHT and USP22 KO Caco-2 cells were subjected to infection with SARS-CoV-2 particles at a MOI of 1. Infected Caco-2 cells were fixed at 24 hours post infection (hpi) and subjected to quantification of SARS-CoV-2 replication via immunofluorescence with the NP antibody recognizing SARS nucleocapsid protein. Interestingly, USP22-deficient cells displayed a prominent decrease in SARS-CoV-2 infection compared to infected WT or NHT Caco-2 cells (Figure 6C), as determined by immunofluorescence of viral protein. In addition, 6 and 24 hpi, SARS-CoV-2-infected USP22-deficient Caco-2 cells had lower genome copy numbers, compared to WT and NHT control cells (Figure 6D). These findings agree with a decreased release of *de novo* infectious SARS-CoV-2 viral particles in supernatants of USP22 KO Caco-2 cells compared to WT and NHT Caco-2 cells (Supplemental Figure 4B). Intriguingly, USP22-STING dKO hIECs exhibit higher SARS-CoV-2 replication rates as well as the formation of more *de novo* infectious viral particles compared to USP22 KO hIECs confirming that the USP22-STING connection also affects antiviral defense against SARS-CoV-2 infection (Figure 6E & F and Supplemental Figure 4C). Together, these findings indicate that USP22 critically controls SARS-CoV-2 infection, replication and the generation of novel infectious viral particles, partially through STING.

## Discussion

Carefully controlled regulation of IFN secretion and signaling is essential for organizing innate immunity, inflammation and anti-viral defense and deregulation of IFNs occur in auto-inflammatory diseases and cancer ^1, 2^. Type I, II and III IFNs elicit complex and intertwined JAK/STAT-based signaling pathways that regulate the expression of IFN stimulated genes (ISG), IFNs, STATs and IRFs with important implications for anti-viral signaling ^9, 10^. Additionally, IFN responsiveness is heavily influenced by IFN receptor affinities, expression and assembly and positive and negative regulation via ISGs, often in cell- and organ-specific manners ^9^. IFN signaling, ISG function and PRR-mediated antiviral defense is carefully controlled by ubiquitination and multiple deubiquitinating enzymes (DUBs) have been linked to the regulation of IFN-specific JAK/STAT pathways and response to viral infection ^38^.

Apart from studying USP22 functions on interferon signaling in mouse models ^60^, previous findings exclusively investigate cellular functions of USP22 upon virus infection and applied overexpression models to investigate USP22 interactions and USP22-mediated ubiquitination ^55, 56^. Here, we are the first to study the basal functions of USP22 in the regulation of ISG expression and STAT signaling in native human intestinal epithelial cells (hIECs). We identify USP22 as negative regulator of type III IFN secretion in basal settings without the addition of exogenous IFNs or by viral infection. Our findings reveal that USP22 regulates both basal and 2’3’-cGAMP-induced STING ubiquitination and activation, even in the absence of ectopic IFNs or viral infection, and loss of STING expression reverses the effects of USP22 KO on IFN signaling. Finally, we test the functional relevance of basal USP22- and STING-mediated IFN and JAK/STAT priming on SARS-CoV-2 infection and identify a critical role of USP22 in the control of SARS-CoV-2 infection, replication and *de novo* formation of infectious viral particles, in a STING-dependent manner.

Despite the finding that USP22 regulates ISG expression in hIECs, USP22 does not exclusively control ISG or IFN-related gene expression. Hence, a large fraction of IFN-unrelated genes is changed while the expression of other genes is not altered upon loss of USP22 expression. Until now, the basis for this selectivity remains unclear. In agreement with previous observations ^50, 72^, loss of USP22 expression in hIECs indeed increased H2Bub1, a hallmark of transcriptionally active chromatin ^73–75^. Interestingly, increased H2Bub1 could also be detected at nucleosomes at ISG-coding genes upon specific deletion of USP22 in the murine hematopoietic system, underlying the upregulation of ISG expression ^60^. This was accompanied by alterations in hematopoietic stem cells (HSCs), myelopoiesis, B cell development, T cell activation, the numbers of B and plasma cells, serum immunoglobulin levels and the appearance of autoantibodies, but not with an increased systemic secretion of IFNs ^60^. This is surprising since IFNs themselves are ISGs as well and IFN expression levels are often maintained at low basal levels to serve as priming signals that allow a fast and adequate increase of IFN responses upon virus infection. Indeed, a large fraction of USP22-regulated ISGs has been demonstrated to be involved as important anti-SARS-CoV-2 countermeasures ^76^. In addition, global and ISG-specific levels of H2Bub1 can be regulated by type I IFN signaling during infection with human adenovirus as well, in a manner depending on human Bre1/RNF20 and the viral E1A protein ^77^. Intriguingly, the RNF20/RNF40 E3 ligase complex, responsible for H2B ubiquitination ^78^, was shown to protect against SARS-CoV-2 infection, and RNF20 becomes cleaved and inactivated by the SARS-CoV-2 protease 3Clpro ^79^. At present, the functional role of 3CLpro-mediated inactivation of RNF20 for H2Bub1 still remains to be addressed.

Since loss of USP22 mostly affects type III IFN expression and secretion, USP22 likely mediates ISG expression both via epigenetic regulatory mechanisms as well as through long-term IFN-mediated priming effects in hIECs. In contrast to type I IFN, type III IFN is mostly sensed in gastro-intestinal and airway epithelia and in the blood-brain barrier ^6–8, 11, 65^. IFN-λ mostly exhibits long-term signaling effects and plays important roles in SARS-CoV-2 infection in airway epithelial and gastro-intestinal cells and organoids and has been shown to critically control antiviral defense ^19, 25–29^.

The susceptibility towards SARS-CoV-2-infections is determined by USP22-mediated regulation of STING. STING is described as mediator of IFN-λ1 production in HT-29 cells, and during viral infection in primary human macrophages in a Ku70-dependent manner ^80, 81^. We furthermore demonstrate for the first time that in the absence of viral infections or exogenous IFN, loss of USP22 expression resulted in basal and 2’3’-cGAMP-induced STING ubiquitination in hIECs. In addition, loss of STING expression decreased the IFN/ISG signaling that occurred under USP22 deficiency, suggesting that STING acts as a physical scaffold for USP22-dependent ubiquitin modifications. STING ubiquitination serves different physiological roles, including determining protein stability, mediating protein-protein interactions and cellular localization ^39–48^. Recently, cGAS-STING activity has emerged as regulator of immunopathology in COVID-19, highlighting the relevant of STING regulation ^82^. STING ubiquitination enables the STING-TBK1 interaction upon cGAS-mediated recognition of cytosolic DNA and is generally associated with activation of ISG expression ^72^. Until now, USP22-mediated STING ubiquitination has only been described upon viral infection and upon ectopic overexpression. For example, overexpressed USP22 modifies ectopically expressed STING with HA-tagged K27 ubiquitin upon HSV-1 infection in HEK293T cells ^56^. USP22 controls nuclear accumulation of IRF3 and type I IFN signaling through KPNA2 deubiquitination only upon infection with SeV and HSV-1 and loss of USP22 expression decreased type I IFN responses upon virus infection, while USP22 deletion in uninfected cells did not trigger basal IFN signaling ^55^. At present, the role of type III IFNs in SeV and HSV-1 infections remains unclear.

Taken together, our findings identify USP22 as central host factor that determines ISG expression and type III IFN production via STING, with important implications for SARS-CoV-2 infection and IFN priming.

## Materials and Methods

### Cell culture, reagents and chemicals

The human colon carcinoma cell line HT-29 was obtained from DSMZ (Braunschweig, Germany) and cultivated in McCoy’s 5A Medium GlutaMAX^TM^-I (Life Technologies, Inc., Eggenstein, Germany), supplemented with 10 % fetal calf serum (FCS) (Biochrom, Ltd., Berlin, Germany) and 1 % penicillin-streptomycin (Invitrogen). The human colorectal adenocarcinoma cell line Caco-2 was provided by Jindrich Cinatl Jr. (Frankfurt am Main, Germany) and maintained in MEM medium (Sigma), supplemented with 10 % FCS, 1 % penicillin-streptomycin, 2 % L-glutamine (Gibco). HEK293T cells were cultivated in Dulbecco’s modified Eagle’s medium (DMEM) supplemented with 10 % FCS and 1 % penicillin/streptomycin. Vero E6 African green monkey kidney cells were obtained from ATCC and maintained in DMEM supplemented with 10 % FCS and 1 % penicillin/streptomycin. Cell lines were cultivated in humidified atmosphere at 37 °C with 5 % CO_2_ and sub-culturing of cells was performed two or three times a week. All cell lines were regularly negatively tested for mycoplasma.

IFN-stimulating DNA (ISD), the cationic lipid-based transfection reagent LyoVec and cyclic [G(2’,5’)pA(3’,5’)p] (2’3’-cGAMP) were obtained from Invivogen (San Diego, USA) and Lipofectamine2000 was obtained from ThermoFisher Scientific (Dreieich, Germany). All other chemicals were obtained from Carl Roth (Karlsruhe, Germany) or Sigma, unless stated otherwise.

### CRISPR/Cas9 gene editing

CRISPR/Cas9 KO cells were generated as described previously ^54^. Briefly, three independent guide RNAs (gRNAs), targeting USP22 (#1: GCCATTGATCTGATGTACGG, #2: CCTCGAACTGCACCATAGGT and #3: ACCTGGTGTGGACCCACGCG), TMEM173 (#1: CATTACAACAACCTGCTACG, #2: GCTGGGACTGCTGTTAAACG, #3: GCAGGCACTCAGCAGAACCA), DDX58 (#1: CATCTTAAAAAATTCCCACA, #2: GGAACAAGTTCAGTGAACTG, #3: TGCATGCTCACTGATAATGA), IFIH1 (#1: CTTGGACATAACAGCAACAT, #2: TGAGTTCCAAAATCTGACAT) or TLR3 (#1: ACGACTGATGCTCCGAAGGG, #2: ACTTACCTTCTGCTTGACAA, #3: GGAAATAAATGGGACCACCA) and control gRNAs (Addgene plasmid #51763, #51762 and #51760) were ligated into pLentiCRISPRv2 (Addgene plasmid # 52961) using restriction cloning. Plasmid fidelity was confirmed using Sanger sequencing. For the generation of viral particles, multiple gene-specific gRNAs were combined and co-transfected with pMD2.G (Addgene plasmid #12259) and psPAX2 (Addgene plasmid #12260) in HEK293T cells using FuGENE HD Transfection Reagent (Promega), according to the manufacturer’s protocol. Viral supernatants were collected 48-and 72-hours post-transfection, pooled and used for transduction in the presence of Polybrene (Sigma-Aldrich), followed by selection with puromycin (Thermo Fischer Scientific). Knockout was confirmed with Western blotting. Where necessary, single-cell clones were selected using limited dilution. Double-knockout (dKO) cells were generated by transduction with USP22-targeting virus first, followed by transduction with viral particles with gRNAs against the appropriate secondary targets and puromycin selection.

### RNA isolation, cDNA synthesis and quantitative real-time PCR

Appropriate cell lines were seeded in 6-well plates (Greiner) 48 hours prior to RNA isolation, treated as indicated or left untreated, followed by extraction of total RNA using the peqGOLD total RNA isolation kit (Peqlab, Erlangen, Germany), according to the manufacturer’s protocol. In brief, cells were lysed in RNA lysis buffer, centrifuged at 12000 x g for 2 min., followed by the addition of an equal volume of 70 % ethanol to the flow-through, after which RNA was bound to RNA-binding columns by centrifugation at 10000 x g for 1 min. Upon washing with RNA Wash Buffer I and two additional wash steps with 80 % ethanol, the column was dried by centrifuging at 12000 x g for 2 min. RNA was eluted with nuclease free water at 12000 x g for 2 min after which 1 µg of RNA was transcribed into cDNA using the RevertAid H Minus First Strand Kit (ThermoFisher Scientific) and random primers, according to the manufacturer’s protocol. Relative mRNA expression levels were quantified using SYBR green-based quantitative real-time PCR (Applied Biosystems, Darmstadt, Germany) using the 7900GR fast real-time PCR system (Applied Biosystems). Data were normalized to 28S housekeeping expression and the relative expression of target gene transcripts levels were calculated compared to the reference transcript using the ΔΔCT method^83^. At least three independent experiments in duplicates are shown. All primers were purchased at Eurofins (Hamburg, Germany). Primer sequences are shown in Supplementary Table 1.

### Gene expression profiling

To quantify global changes in gene expression, RNA was isolated as described above, followed by a DNase digest upon RNA binding using the peqGOLD DNase Digest Kit, according to the manufacturer’s instructions. Samples were processed and gene expression was profiled at the DKFZ Genomics and Proteomics Core Facility (Heidelberg, Germany) using the Affymetrix human Clariom S array.

### Gene expression profiling analysis

Raw .CEL files were processed with the oligo R package ^84^ and the normalized intensities were obtained after RMA normalization. Genes with differential expression between NHT control and USP22 KO have been identified using the linear model-based approach limma R package ^85^. An adjusted P-value < 0.05 was considered significant. Gene-set enrichment analysis was performed with gage R package ^86^ using the MSigDB ^87^ as gene set repository. An adjusted P-value < 0.05 was considered significant.

### Multiplex quantification of cytokine secretion

Cells were seeded in 2 ml cell culture medium and supernatant was collected after 66 h, centrifuged at 300 x g, 4 °C for 5 min. and frozen in liquid nitrogen. Samples were analyzed using the LEGENDplex™ Human Anti-Virus Response Panel multiplex assay (BioLegend, San Diego, CA, USA) following the manufacturer’s protocol. The analysis was performed with the BD FACSVerse™ flow cytometer (BD Biosciences, San Jose, CA, USA). At least 300 events were acquired per analyte. The data was analyzed with the LEGENDplex v.8 software (BioLegend).

### Western Blot analysis

The indicated cell lines were seeded two days before lysis and treated as indicated, or left untreated. Lysis was done on ice using RIPA lysis buffer (50 mM Tris–HCl pH 8, 150 mM NaCl, 1 % Nonidet P-40 (NP-40), 150 mM MgCl_2_, 0.5 % sodium deoxycholate), with phosphatase inhibitors (1 mM sodium orthovanadate, 1 mM β-glycerophosphate, 5 mM sodium fluoride), protease inhibitor cocktail (Roche, Grenzach, Germany), 0.1 % sodium dodecyl sulfate (SDS) and Pierce Universal Nuclease (Thermo Fisher Scientific) for 30 min, followed by centrifugation at 18000 x g for 25 min. at 4 °C. Protein concentrations of the cell lysates were measured using the BCA Protein Assay Kit from Pierce™, according to the manufacturer’s instructions. For Western blot detection, 20-40 µg of the lysates were boiled in Laemmli loading buffer (6x Laemmli: 360 nM Tris Base pH 6.8, 30 % glycerol, 120 mg/ml SDS, 93 mg/ml dithiothreitol (DTT), 12 mg/ml bromophenol blue) at 95 °C for 5 min, followed by Western blot analysis. The following antibodies are used: rabbit anti-STING (13647S, Cell Signaling Beverly, MA, USA), rabbit anti-phospho-STAT1 (9167L, Cell Signaling), mouse anti-STAT1 (9176S, Cell signaling), rabbit anti-USP22 (#ab195298, Abcam), mouse anti-glyceraldehyde 3-phosphate dehydrogenase (GAPDH) (5G4cc, HyTest, Turku, Finland), mouse anti-Vinculin (#V9131-100UL, Merck), rabbit anti-TBK1 (ab40676, Abcam), rabbit anti-phospho-TBK1 (ab109272, Abcam), rabbit anti-Histone H2B (#07-371, Merck), mouse anti-Ubiquityl-Histone H2B (#05-1312, Merck), rabbit anti-p65 (sc-372X, Santa Cruz Biotechnologies, Santa Cruz, CA, USA), rabbit anti-phospho-p65 (3033S, Cell Signaling), mouse anti-IRF3 (sc-33641, Santa Cruz), rabbit anti-phospho-IRF3 (4947S, Cell Signaling), rabbit anti-RIG-I (3743S, Cell Signaling), rabbit anti-MDA5 (5321S, Cell Signaling), rabbit anti-TLR3 (6961S, Cell Signaling), mouse anti-ISG56 (PA3-848, Thermo scientific), rabbit anti-MX1 (37849S, Cell Signaling), rabbit anti-IRF9 (76684S, Cell Signaling), rabbit anti-ISG20 (PA5-30073, Thermo scientific), rabbit anti-γ-H2AX (phospho Ser139) (NB100-384, Novus Biologicals) and mouse anti-NF-κB p52 (05-361, Millipore). Secondary antibodies labeled with horseradish peroxidase (HRP) were used for detection with enhanced chemiluminescence (Amersham Bioscience, Freiburg, Germany). HRP-conjugated goat anti-mouse IgG (ab6789, Abcam) was diluted 1:10000 and HRP-conjugated goat anti-rabbit IgG (ab6721, Abcam) was diluted 1:30000 in 5 % milk powder in PBS with 0.2 % Tween 20 (PBS-T). When necessary, membranes were stripped using 0.4 M NaOH for 10 min, followed by 1 h of blocking and incubation with a second primary antibody. Representative blots of at least two independent experiments are shown. When detected on separate membranes, only one representative loading control is shown for clarity.

### Stimulation of STING with 2’3’-cGAMP

The indicated cell lines were seeded 24 or 48 hours prior to stimulation in P/S-free cell culture medium. For stimulation, culture medium was removed and cell lines were permeabilized by incubation with digitonin buffer (50 mM HEPES, 100 mM KCl, 3 mM MgCl_2_, 0.1 mM dithiothreitol, 85 mM sucrose, 0.2 % bovine serum albumin, 1 mM ATP, 5 µg/ml Digitonin) pH 7 in the presence or absence of 10 µg/ml 2’3’-cGAMP for 10 min. at 37 °C. After incubation, the permeabilization buffer was replaced with P/S-free cell culture medium and further incubated at 37 °C/5 % CO_2_ for the indicated time points.

### PRR stimulation with poly(I:C) and ISD

The indicated HT-29 cells were seeded 24 hours prior to treatment in sterile 6-well plates (Greiner). For each well, two µg of ISD (Invivogen) were pre-mixed with OptiMEM and, after 5 min. incubation at room temperature, mixed with premixed Lipofectamine2000-OptiMEM at a ratio of 3:1, according to the manufacturer’s instructions. After incubation for 15 min. at room temperature, the indicated transfection mixes were added to the cells in P/S free medium. Cell lysis with RIPA or RNA lysis buffer was performed after 24 h. For stimulation with poly(I:C), the indicated HT-29 cells were seeded as described above and for each well, 2 µg of poly(I:C) was mixed with 20 µl LyoVec (Invivogen), incubated for 15 min. at room temperature to allow the formation of lipid-RNA complexes. The transfection mix was then added to the indicated HT-29 cells in P/S free medium at a 1:20 volume ratio and incubated for 24 h, after which cells were processed for Western blot or RNA isolation.

### Tandem Ubiquitin Binding Entity (TUBE) pull-down analysis

Ubiquitinated proteins were enriched using GST-tagged tandem-repeated ubiquitin-binding entities (TUBEs) ^88^, as described before ^54^. Briefly, the indicated cell lines were seeded 48 hours prior to lysis and/or treatment, harvested in NP-40 lysis buffer (50 mM NaCl, 20 mM Tris pH 7.5, 1 % NP-40, 5 mM EDTA, 10 % Glycerol) supplemented with 25 mM NEM, 1 mM sodium orthovanadate, 1 mM sodium fluoride, 0.5 mM phenylmethylsulfonyl fluoride, protease inhibitor cocktail and Pierce Universal Nuclease on ice for 30 min. GST-TUBE beads were washed once with NP-40 buffer an incubated with 3 mg of protein lysate over night at 4 °C. Beads were washed four times with NP-40 buffer, followed by elution of ubiquitinated proteins by boiling in 2x Laemmli loading buffer at 96 °C for 6 min. Ubiquitinated proteins were analyzed using Western blot analysis.

### SARS-CoV-2 infection

SARS-CoV-2 (strain BavPat1/2020) was obtained from the European Virology Archive and amplified in Vero E6 cells and used at passage 3. Virus titers were determined by TCID50 assay. Caco-2 cells were infected using a MOI of 1 virus particle per cell. Medium was removed from Caco-2 cells and virus was added to cells for 1 h at 37°C. Viral supernatants were removed, infected cells were washed once with PBS and media was added back to the cells. Virus infection was monitored 24 h post-infection.

### TCID50 virus titration

Vero E6 cells were seeded (20000 per well) in 96-well plates 24 h prior to infection. A volume of 100 µl of viral supernatant from the indicated SARS-CoV-2-infected Caco-2 cells was added to the first well. Seven 1:10 dilutions were made (all samples were performed in triplicate). Infections were allowed to proceed for 24 h. At 24 h post infection (hpi), cells were fixed in 2 % paraformaldehyde (PFA) for 20 minutes at room temperature. PFA was removed and cells were washed twice in PBS and permeabilized for 10 min. at room temperature in 0,5 % Triton-X/PBS. Cells were blocked in a 1:2 dilution of LI-COR blocking buffer (LI-COR, Lincoln, NE, USA) for 30 min at room temperature. Infected cells were stained with 1:1000 diluted anti-dsRNA (J2) for 1 h at room temperature, washed three times with 0.1 % PBT-T, followed by incubation with secondary antibody (anti-mouse CW800) and DNA dye Draq5 (Abcam, Cambridge, UK), diluted 1:10000 in blocking buffer and incubated for 1 h at room temperature. Cells were washed three times with 0.1 % PBS-T and imaged in PBS on a LI-COR imager.

### Quantification of viral RNA

At 24 hpi, RNA was extracted from infected or mock-treated Caco-2 cells using the Qiagen RNAeasy Plus Extraction Kit (Qiagen, Hilden, Germany). For quantifying the SARS-CoV-2 genome abundance in mock and infected samples, cDNA was generated using 250 ng of RNA with the iSCRIPT reverse transcriptase (BioRad, Hercules, CA, USA), according to the manufacturer’s instructions. qRT-PCR was performed using iTaq SYBR green (BioRad) following the instructions of the manufacturer and normalized on TBP. Primers were ordered at Eurofins, Luxemburg and are listed in Supplementary Table 1.

### Indirect Immunofluorescence Assay

Cells seeded on iBIDI glass bottom 8-well chamber slides. At 24 post-infection, cells were fixed in 4% paraformaldehyde (PFA) for 20 mins at room temperature (RT). Cells were washed and permeabilized in 0.5% Triton-X for 15 mins at RT. Primary antibody SARS-CoV NP (Sino biologicals MM05) were diluted in phosphate-buffered saline (PBS) and incubated for 1h at RT. Cells were washed in 1X PBS three times and incubated with secondary antibodies goat-anti mouse Alexa Fluor 568 and DAPI for 45 mins at RT. Cells were washed in 1X PBS three times and maintained in PBS. Cells were imaged by epifluorescence on a Nikon Eclipse Ti-S (Nikon).

### Statistical analysis

Significance was assessed using Student’s t-test (two-tailed distribution, two-sample, equal variance) using Microsoft Excel, unless indicated otherwise. *P*-values < 0.05 are considered significant (* *P* < 0.05; ** *P* < 0.01; *** *P* < 0.001, n.s.: not significant).

### Resource availability

Further information and requests for resources and reagents should be directed to and will be fulfilled by the corresponding author, Sjoerd J. L. van Wijk.

### Materials availability

All unique reagents generated in this study are available from the corresponding author without restriction.

### Data and code availability

Microarray data are available on Gene Expression Omnibus under the accession number GSE190036.

## Acknowledgments

The authors thank the members of the van Wijk lab for advice, discussions and support during the study, Dr. M. Bewerunge-Hudler and her team from the Genomics and Proteomics Core Facility, German Cancer Research Center/DKFZ, Heidelberg, Germany for help and support with performing the microarray analysis and Christina Hugenberg for proofreading. S.J.L.v.W. is supported by the Deutsche Forschungsgemeinschaft (DFG) (WI 5171/1-1, FU 436/20-1 and project-ID 259130777 – SFB 1177), the Deutsche Krebshilfe (70113680), the Frankfurter Stiftung für krebskranke Kinder and the Dr. Eberhard and Hilde Rüdiger Foundation. M.B. is supported by the DFG – CRC 850 subprojects C9 and Z1, CRC1479 (Project ID: 441891347-S1), CRC 1160 (Project Z02), CRC1453 (Project ID 431984000 -S1) and TRR167 (Project Z01), the German Federal Ministry of Education and Research by MIRACUM within the Medical Informatics Funding Scheme (FKZ 01ZZ1801B). S.B. was supported by DFG project numbers 415089553 (Heisenberg program), 240245660 (SFB1129), 278001972 (TRR186), and 272983813 (TRR179), the state of Baden-Württemberg (AZ: 33.7533.-6-21/5/1), the BMBF (01KI20198A) and within the Network University Medicine - Organo-Strat COVID-19. M.L.S. was supported by the BMBF (01KI20239B) and DFG project 416072091.

## Author contributions

R.K. performed experiments and analyzed data with help from J.R., S.S. and L.K., M.L.S and S.B. performed SARS-CoV-2 infections and accompanying experiments, gene expression analysis was performed by G.A. and M.B., R.S. provided access and support with the LEGENDplex analysis. D.B and J.C.Jr. provided the Caco-2 cell line and expertise. R.K. and S.J.L.v.W. conceived the project and wrote the manuscript. All authors have read, commented and agreed on the submitted version of the manuscript.

## Supplementary Information

**Summary:** Four Supplementary Figures including Supplementary Figure legends and a Supplementary Table

**Supplemental Figure 1 (related to Figure 1).**
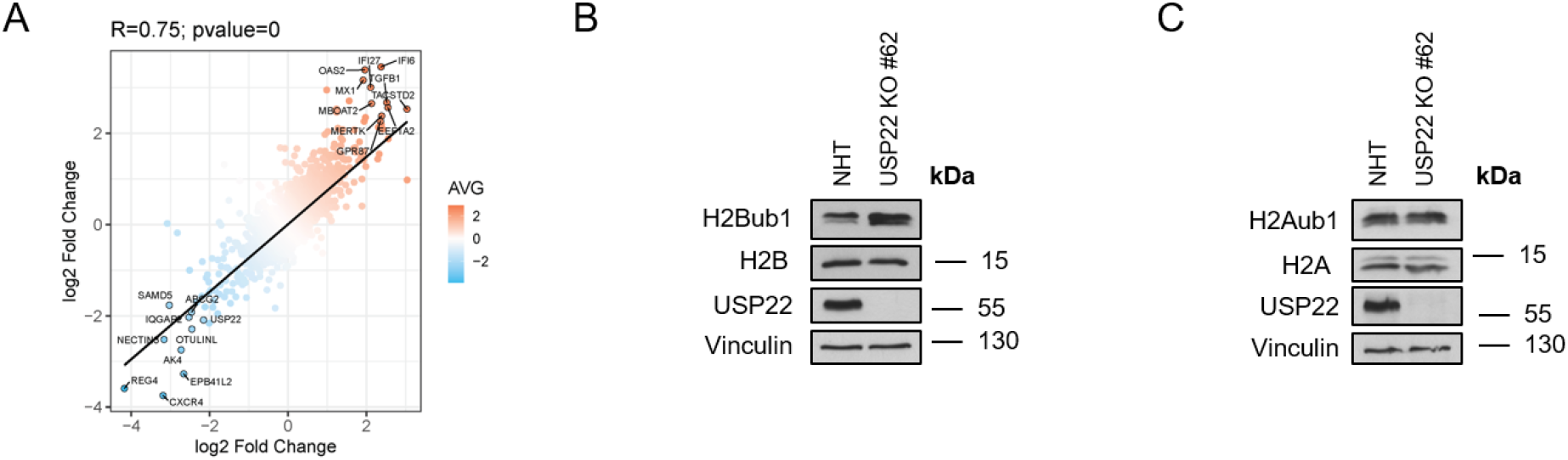
**A.** Scatter plot demonstrating the changes in gene expression of CRISPR/Cas9 control (NHT) HT-29 cells with two independent single-cell HT-29 USP22 KO clones (#16 and #62). Color code represents the average log2 foldchange. **B, C.** Western blot analysis of mono-ubiquitinated (H2Bub1) and total levels of Histone 2B (H2B) (**B**) and Histone 2A (H2A) (**C**) as well as USP22 in control and USP22 KO HT-29 cells (USP22 KO #62). Vinculin served as loading control. Representative blots of at least two different independent experiments are shown.

**Supplemental Figure 2 (related to Figure 3).**
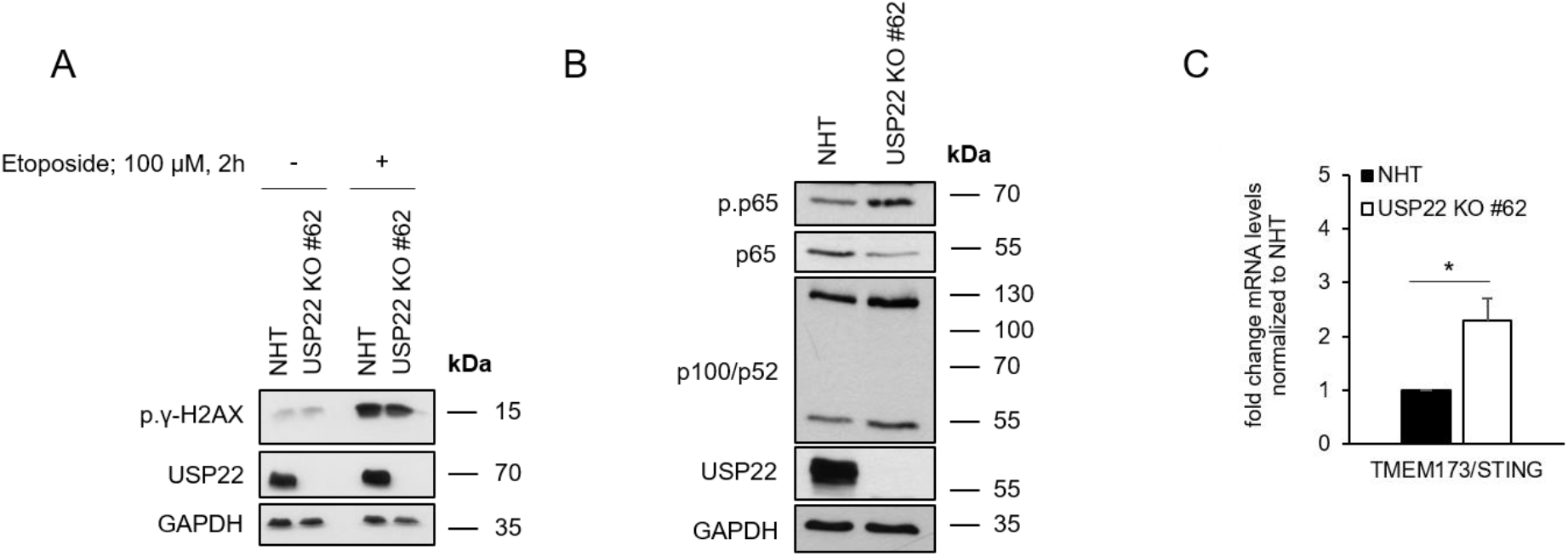
**A.** Western blot analysis of phosphorylated γ-H2AX (p.γ-H2AX) and USP22 expression levels in control (non-human target: NHT) and CRISPR/Cas9-generated USP22 knock-out (KO) HT-29 cells (USP22 KO) subjected to vehicle or etoposide (100 µM) for 2 h. GAPDH served as loading control. Representative blots of at least two different independent experiments are shown. **B.** Western blot analysis of phosphorylated and total p65, p100/p52 and USP22 expression levels in control and USP22 KO HT-29 cells (USP22 KO #62). GAPDH served as loading control. Representative blots of at least two different independent experiments are shown. **C.** Basal mRNA expression levels of TMEM173/STING in control and USP22 KO HT-29 cells (USP22 KO #62) using qRT-PCR. Gene expression was normalized against 28S mRNA and is presented as x-fold mRNA expression compared to NHT. Mean and SD of three independent experiments in triplicate are shown. *P < 0.05.

**Supplemental Figure 3 (related to Figure 5).**
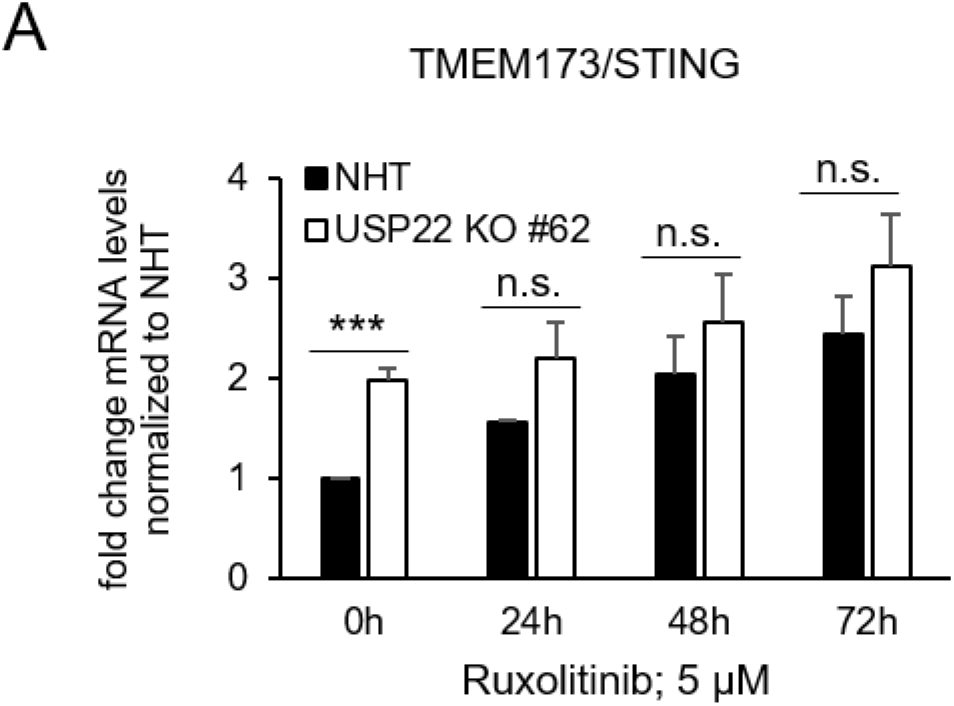
**A.** mRNA expression levels of TMEM173/STING in control (non-human target: NHT) and CRISPR/Cas9-generated USP22 knock-out (KO) HT-29 cells (USP22 KO) using qRT-PCR. Cells were treated with ruxolitinib (5 µM) for the indicated timepoints. Gene expression was normalized against 28S mRNA and is presented as x-fold mRNA expression compared to NHT. Mean and SD of three independent experiments in triplicate are shown. ***P < 0.001, n.s. not significant.

**Supplemental Figure 4 (related to Figure 6).**
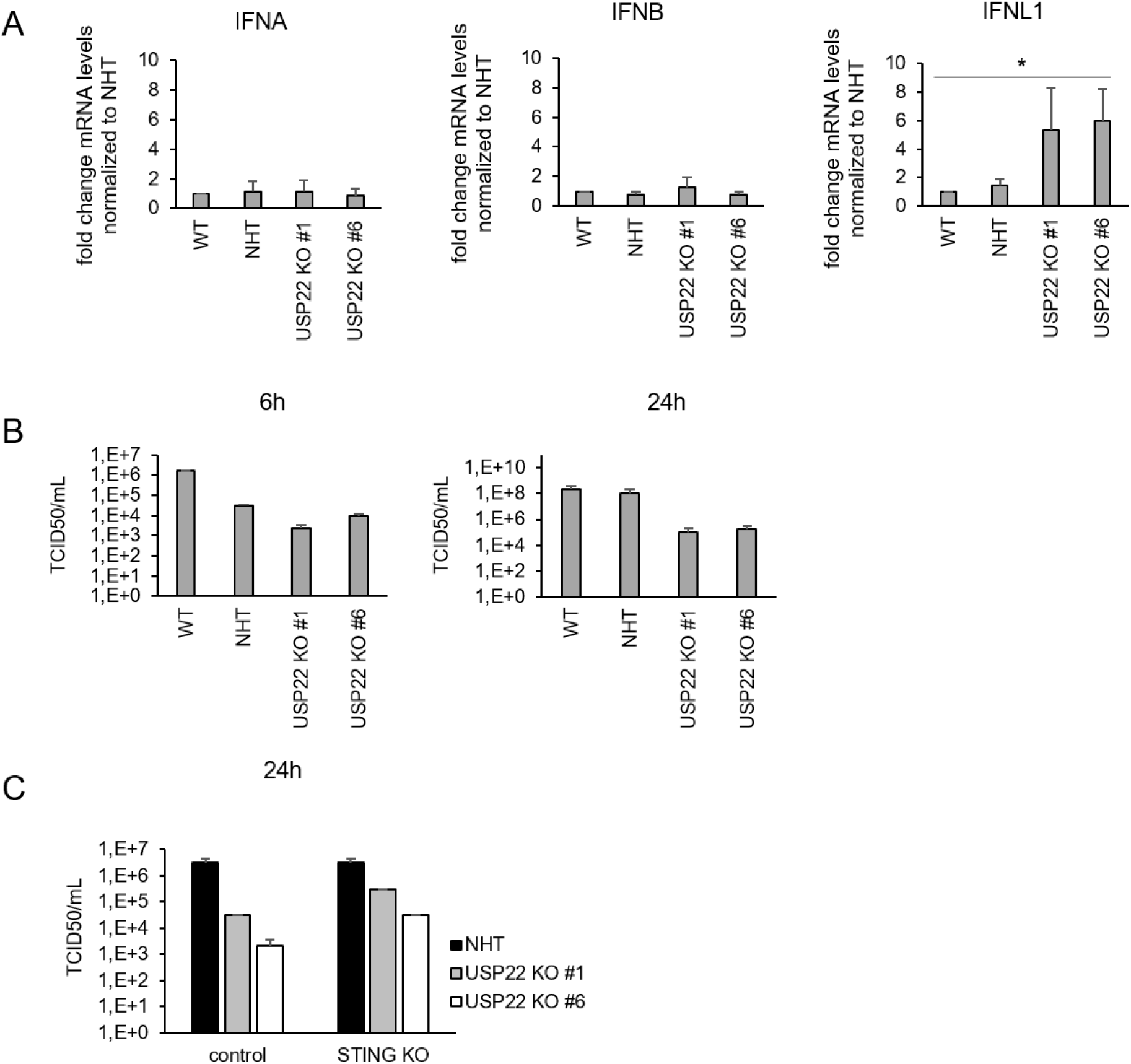
**A.** Basal mRNA expression levels of IFNA, IFNB and IFNL1 in wild-type (WT), control (non-human target: NHT) and CRISPR/Cas9-generated USP22 knock-out (KO) Caco-2 single clones (USP22 KO #1 and #6). Gene expression was normalized against 28S mRNA and is presented as x-fold mRNA expression compared to NHT. Mean and SD of four (IFNA, IFNB) or three (INFL1) independent experiments in triplicate are shown. *P < 0.05. **B.** TCID50/mL, determined via titration of supernatant from SARS-CoV-2-infected WT, control and USP22 KO Caco-2 cells (USP22 KO #1 and #6) 6 and 24 hpi on Vero cells. Mean and SD of three independent experiments in triplicate are shown. **C.** Idem, 24 hpi, supernatant additionally of NHT-, USP22 KO #1-and #6-STING dKO Caco-2 cells.

**Supplementary Table 1:**
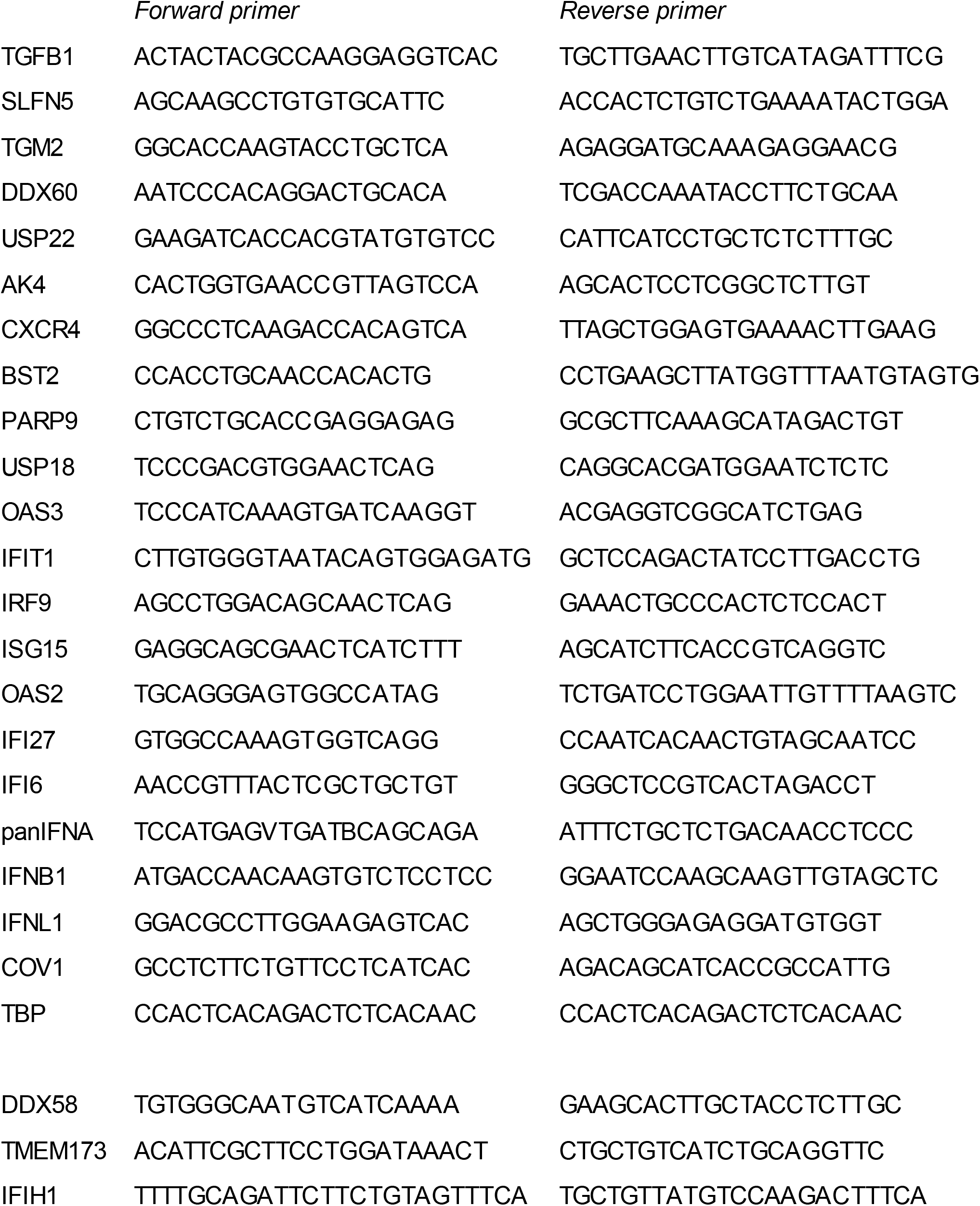
List of qRT-PCR primers used in this study.

